## Supplementary File (Figures S1-6) for "Direct inference and control of genetic population structure from RNA sequencing data"

**Supplementary Materials**

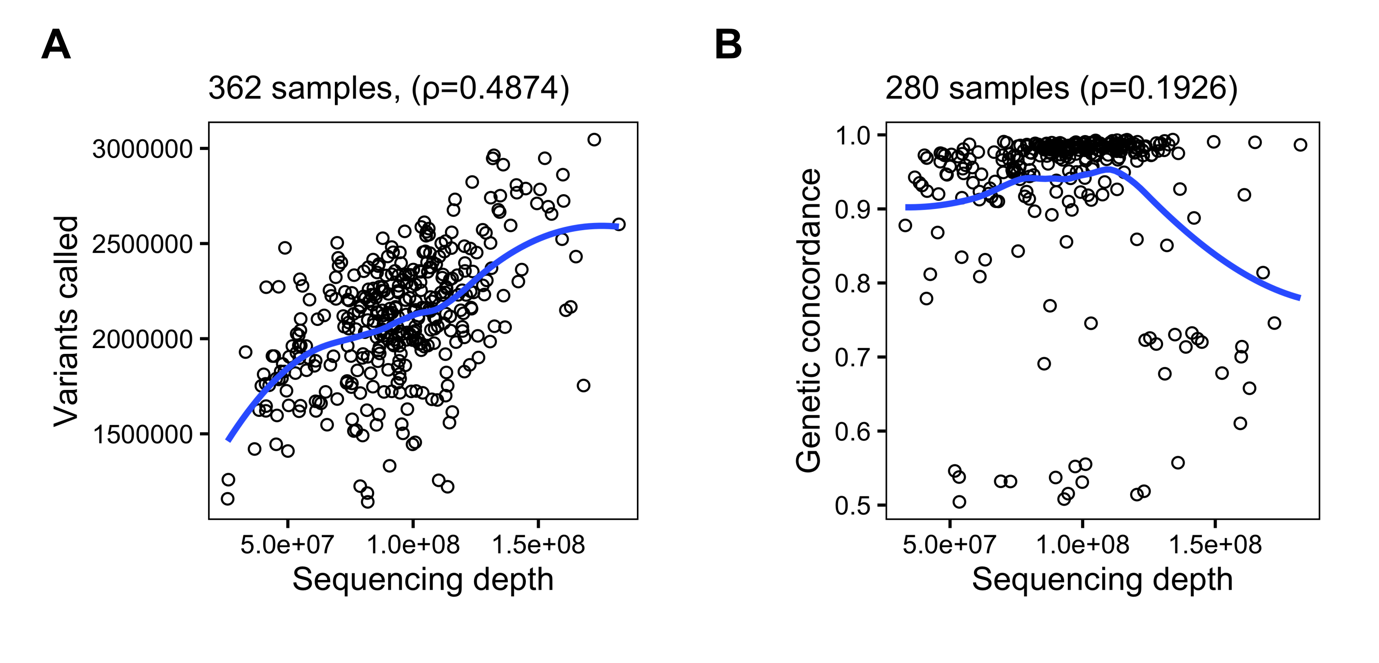

**Figure S1. (A)** Raw sequencing depth correlated moderately to total variants called per sample. **(B)** When looking at only samples found in both RNAseq and array SNP sets (280 samples), weaker Spearman correlation was found between sequencing depth and the genetic concordance level of RNAseq and array SNPs.

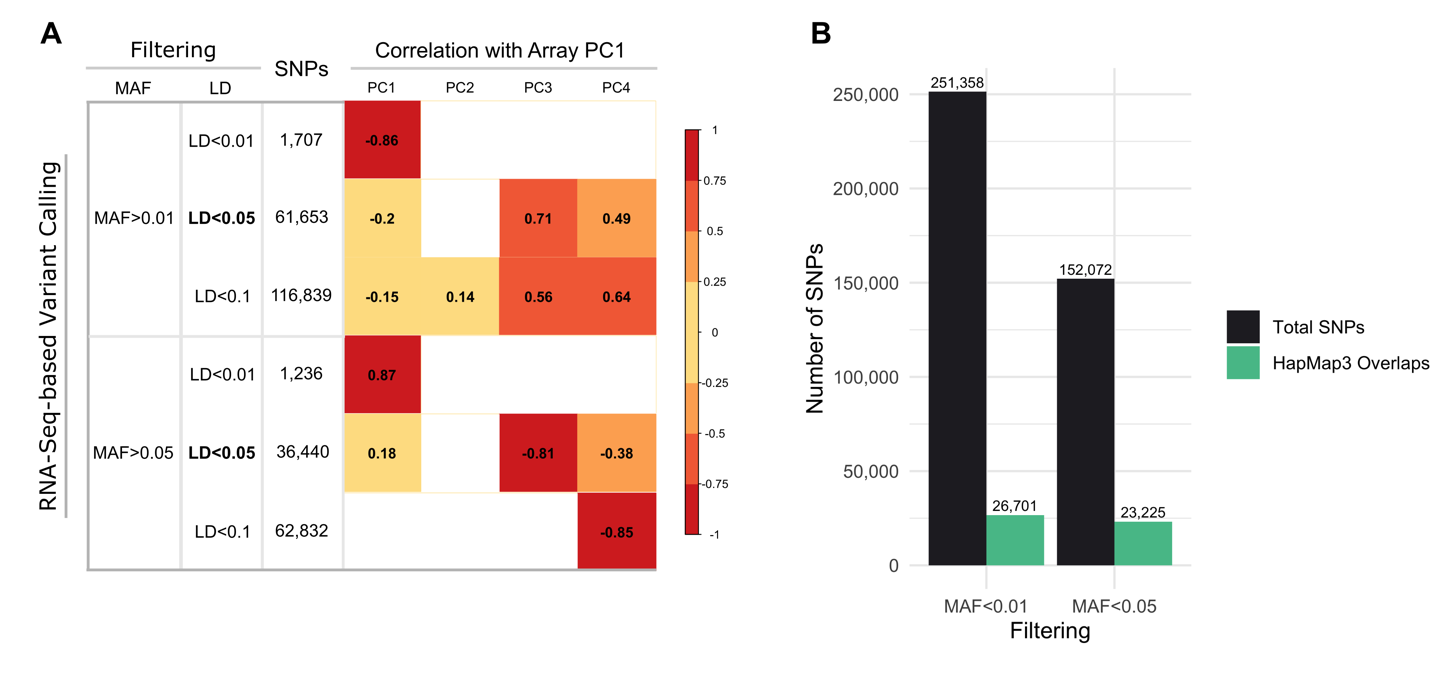

**Figure S2.** After initial removal of duplicated, palindromic, and non-autosomal variants, further filtering of variants from RNAseq data is necessary in acquiring a healthy number of high-quality SNPs, which could be done based on minor allele frequency (MAF), overlapping with a known list of high-quality SNPs such as HapMap3, as well as linkage disequilibrium (LD)-thinning. **(A)** Six different filters based on two different MAF and three different LD requirements were tested, from which RNAseq-based genetic PCs (RG-PCs) were calculated and samples with paired genotyped arrays were compared to the array-based genetic PCs (only PC1 from array was shown, as it represents the main stratification within the population). While attempts using LD<0.01 showed the highest absolute correlation with the array PC1 (average |r|=0.865), the numbers of SNPs left only averaged 1,471. The attempt with MAF>0.05 and LD<0.1 had comparable absolute correlation (|r|=0.85), yet it was achieved on PC4 with no significant correlations on the three previous PCs. Attempts using LD<0.05 had the best balance between number of PCs as well as decent initial correlations with array PC1 without overlapping with HapMap3 SNP list. **(B)** When overlapping with SNPS from HapMap3, the attempt using MAF>0.05 returned 23,225 overlapping SNPs from over 152,000 initial SNPs. With over 251,000 initial SNPs, filtering with MAF>0.01 returned 24,424 overlaps with HapMap3. Since LD filter is necessary after subsetting, filtering with MAF>0.05 (and eventually LD<0.05 after getting overlaps) is preferable in achieving high-quality SNPs without losing too much information.

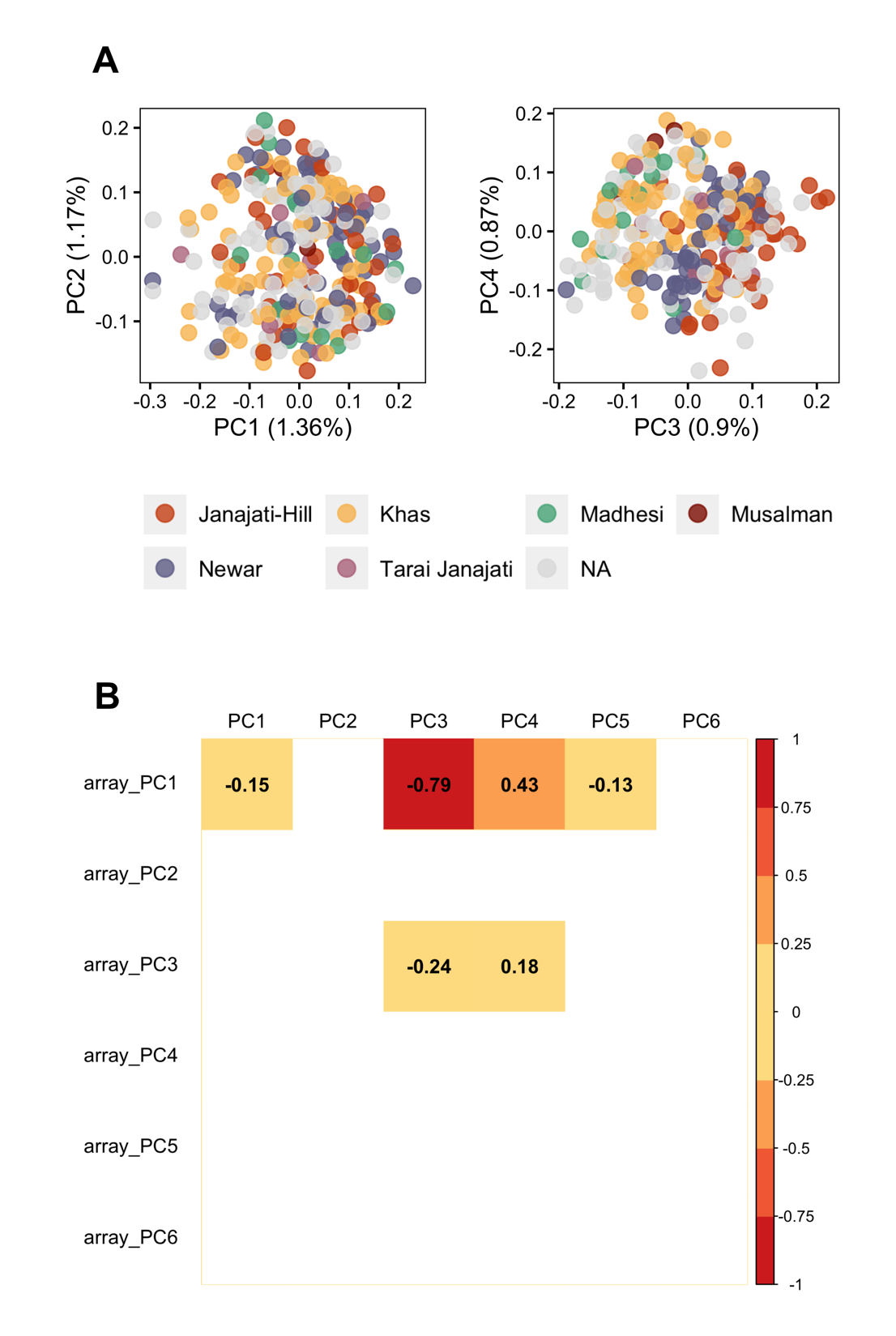

**Figure S3.** **(A)** PCA plots based on SNPs from RNAseq variant calling after MAF and LD filtering (MAF>0.05, LD<0.05) but without subsetting to HapMap3 SNPs. Population structure based on self-reported ethnicity is not clearly visible in principal components (PCs) 1-2. This is supported by **(B)** the correlations between the first 6 PCs of array SNPs and the first 6 RNAseq-based genetic principal components above, as the population structure could only be inferred starting from PC3, suggesting that the main PCs 1-2 captured considerable variations unrelated to the population structure. The correlation map only shows significant Spearman correlations (correlations with p-value>0.05 were included as blanks).

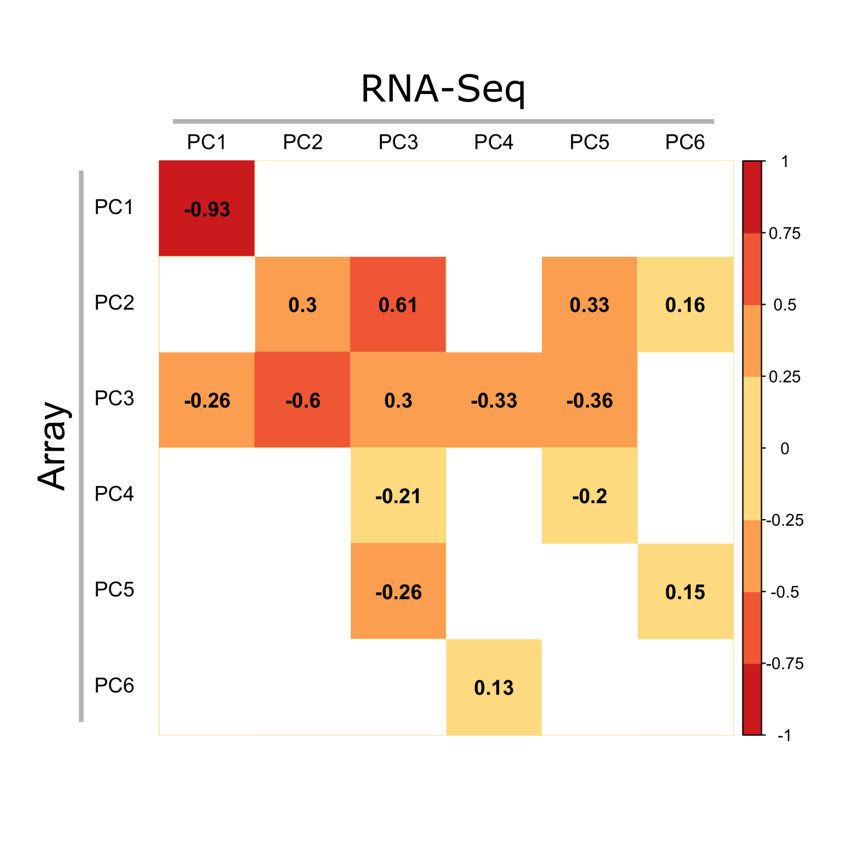

**Figure S4.** Correlation map between the first 6 principal components (PCs) of array SNPs and the first 6 RNAseq-based genetic principal components (RG-PCs, after subsetting with HapMap3 SNP list), showing only significant Spearman correlations (correlations with p-value>0.05 were included as blanks).

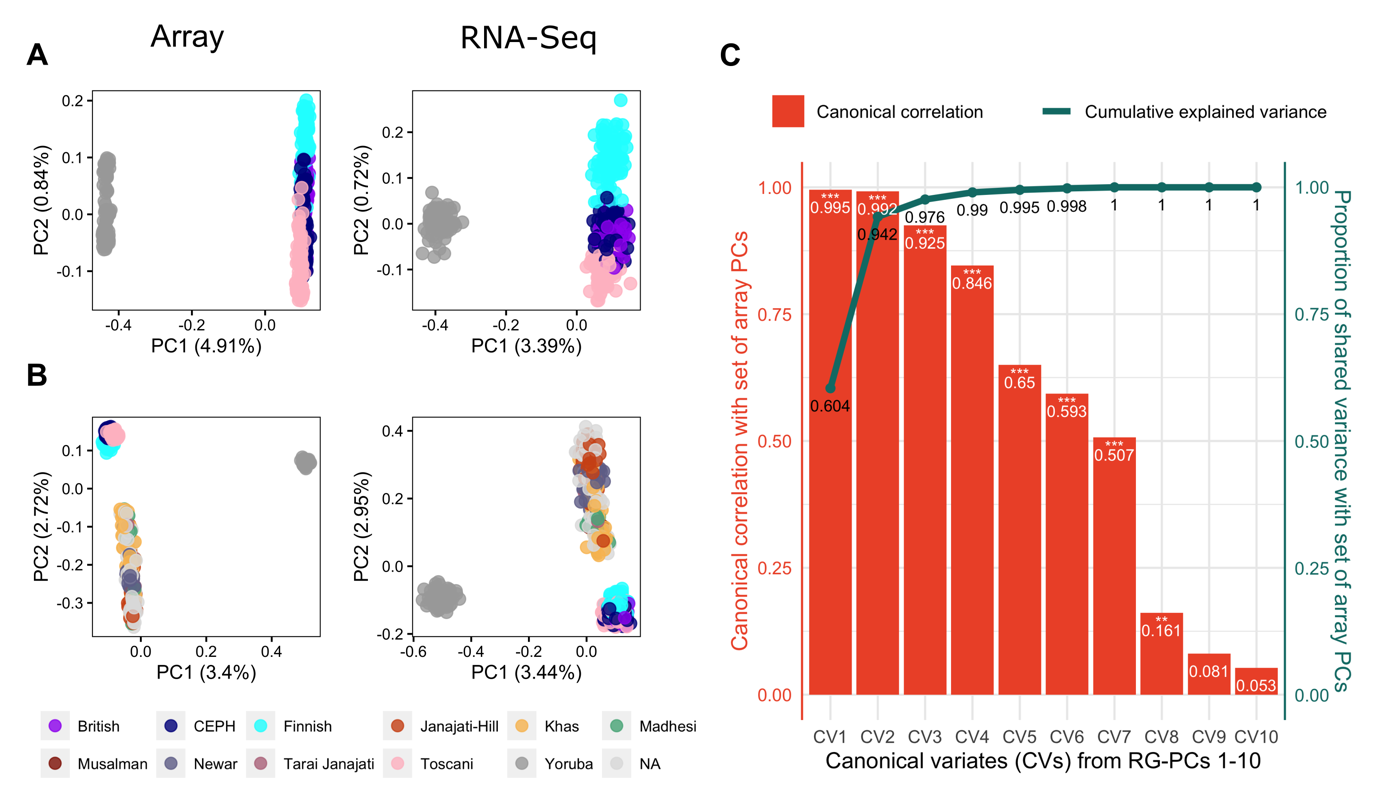

**Figure S5.** RGStraP was able to capture population structure from RNAseq comparable to paired array genotypes, as seen in the PCA plots when analyzing **(A)** Geuvadis samples, as well as **(B)** both Nepal and Geuvadis samples. RGstraP was able to capture the variations between and within populations. (C) Canonical correlation analysis between ten RG-PCs and ten array PCs of both the Geuvadis and Nepal samples showed significant (Wilks’ Lambda, p-value<0.05) correlations for the first 8 canonical variates (CVs) between the two sets. The first 4 CVs from 10 RG-PCs strongly captured the genetic information from array PCs (R_c1_=0.995, R_c2_=0.992, R_c3_=0.925, R_c4_=0.846). The cumulative proportion of shared variance between the two sets reached 0.976 from just the 3 CVs.

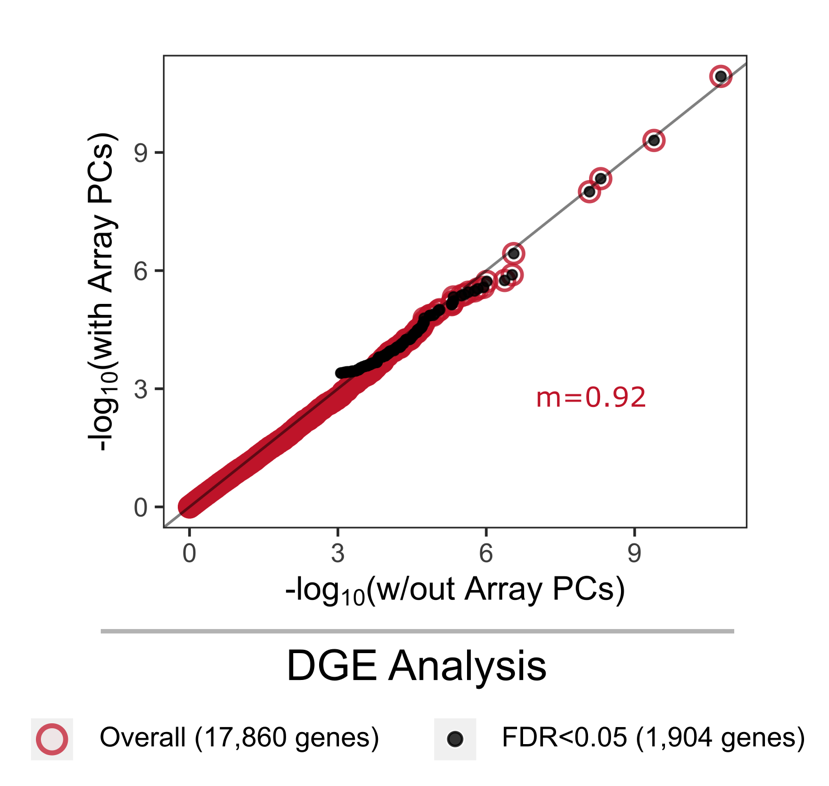

**Figure S6.** Quantile-quantile (Q-Q) plots showing the distribution of the probabilities between analyses with and without including genetic PCs. Differential gene expression analysis results between samples of different sex show systematic reduction in test statistics when including array-based genetic PCs as covariates compared to without, demonstrated by the low systematic inflation metric (m).
